# Ori-Finder 2022: A Comprehensive Web Server for Prediction and Analysis of Bacterial Replication Origins

**DOI:** 10.1101/2022.08.09.503306

**Authors:** Meijing Dong, Hao Luo, Feng Gao

**Author notes:** Corresponding author. (Gao F). Equal contribution (Meijing Dong, Hao Luo).

## Abstract

The replication of DNA is a complex biological process that is essential for life. Bacterial DNA replication is initiated at genomic loci referred to as replication origins (*oriC*s). Integrating the Z-curve method, DnaA box distribution, and comparative genomic analysis, we developed a web server to predict bacterial *oriC*s in 2008 called Ori-Finder, which contributes to clarify the characteristics of bacterial *oriC*s. The *oriC*s of hundreds of sequenced bacterial genomes have been annotated in their genome reports using Ori-Finder and the predicted results have been deposited in DoriC, a manually curated database of *oriC*s. This has facilitated large-scale data mining of functional elements in *oriC*s and strand-biased analysis. Here, we describe Ori-Finder 2022 with updated prediction framework, interactive visualization module, new analysis module, and user-friendly interface. More species-specific indicator genes and functional elements of *oriC*s are integrated into the updated framework, which has also been redesigned to predict *oriC*s in draft genomes. The interactive visualization module displays more genomic information related to *oriC*s and their functional elements. The analysis module includes regulatory protein annotation, repeat sequence discovery, homologous *oriC* search, and strand-biased analyses. The redesigned interface provides additional customization options for *oriC* prediction. Ori-Finder 2022 is freely available at http://tubic.tju.edu.cn/Ori-Finder2022 and https://tubic.org/Ori-Finder2022.

## Introduction

As a complex and essential process of cell life, DNA replication is strictly regulated to ensure the accurate transfer of genetic material from parents to offspring. Identification and characterization of replication origins (*oriC*s) can provide new insights into the mechanisms of DNA replication as well as cell cycle regulation and facilitate drug development [1], genome design [2], plasmid construction [3] etc. Therefore, various experimental approaches such as two-dimensional agarose gel electrophoresis [4], assay of autonomously replicating sequence activity [5], and marker frequency analysis (MFA) [6] have been developed to identify bacterial *oriC*s. Microarray based whole-genome MFA [7] as well as high-throughput sequencing-based MFA [8] with higher resolution have been proposed to generate the replication maps of genomes and to locate *oriC*s. Detecting interactions between DNA and proteins can also provide evidence for predicted *oriC*s [9].

However, the rapid accumulation of sequenced genomes has rendered identifying *oriC*s in all of them impossible using experimental methods. Therefore, the development of bioinformatics algorithms to predict *oriC*s on a large scale is particularly important. Classical *in silico* methods, such as GC skew [10], cumulative GC skew [11], and oligomer skew [12] have been proposed based on DNA asymmetry. Furthermore, Oriloc was developed to predict bacterial *oriC*s by analyzing local and systematic deviations of base composition within each strand [13]. However, these methods only provided the approximate location without the precise boundary of predicted *oriC*s. In addition, they cannot accurately predict *oriC*s in bacterial genomes without a typical GC skew, which is universal sometimes for genomes in certain phylum, such as Cyanobacteria [14, 15]. Although DNA asymmetry is the most common characteristic used for predicting *oriC*s, Mackiewicz *et al*. also found that the prediction could be improved more by considering *dnaA* and DnaA box clusters [16]. However, DnaA box motifs are often species-specific, and the *oriC* is not always close to the *dnaA* gene in some species. Considering these factors, the Ori-Finder web server was developed to provide users with a more convenient and accurate tool for predicting *oriC*s [17].

Since it was introduced in 2008, Ori-Finder has been widely used to help investigators identify *oriC*s. To date, Ori-Finder has been used to identify *oriC*s in hundreds of sequenced bacterial genomes in their genome reports [18-20], and dozens of the predicted *oriC*s have been experimentally confirmed [21-24]. Furthermore, Ori-Finder predictions have led to new discoveries. For example, each bacterial chromosome is generally considered to carry a single *oriC*. However, Ori-Finder predictions indicate that multiple *oriC*s may occur on a bacterial chromosome [25, 26], and this opinion has been used to explain the experimental results of investigations into single *Achromatium* cells [27]. Naturally occurring single chromosome in *Vibrio cholerae* strain harbor two functional *oriC*s, which provides strong support for our opinion [28]. Ori-Finder provides a large number of *oriC*s as resources for data mining. Particularly, the *oriC*s identified by Ori-Finder, including those confirmed by experiments *in vivo* and *in vitro*, have been organized into the DoriC database [29-31] available at http://tubic.org/doric. Therefore, the data for *oriC* characteristics can be mined on a large scale [32, 33]. For example, vast amounts of *oriC* data can be used to identify and analyze functional elements, such as DnaA boxes and DnaA-trios [34, 35]. Finally, Ori-Finder facilitates analyses of strand-biased biological characteristics that are closely associated with DNA replication, transcription, and other biological processes [10, 36]. The Ori-Finder web server and DoriC database have been extensively applied to in strand-biased analyses, such as base composition [37, 38], gene orientation [39], and codon usage [40]. Ori-Finder has also been recommended as a software tool to identify replichores [41].

Bacterial *oriC*s generally contain several functional elements, such as DnaA-binding sites, AT-rich DNA unwinding elements (DUEs), and binding sites for proteins that regulate replication initiation [42]. These functional elements play important roles in the initiation of DNA replication, which should be considered in the prediction of *oriC*s. Most of bacterial *oriC*s contain DnaA box clusters that are recognized and bound by DnaA proteins. Therefore, the DnaA box cluster is considered as an important characteristic for predicting *oriC*s [16]. DnaA box is usually a 9-bp non-palindromic motif, such as the perfect *Escherichia coli* DnaA box TTATCCACA. Species-specific DnaA box motifs, such as TTTTCCACA in Cyanobacteria and AAACCTACCACC in *Thermotoga maritima* have been identified [43]. In addition, degenerated DnaA boxes have also been identified within *oriC*s in some species, such as 6mer ATP-DnaA boxes (AGATCT) in *E. coli* [44]. Although degenerate DnaA boxes can also bind DnaA protein, only the broadly conserved DnaA box is considered for *oriC* prediction here.

The DnaA protein not only interacts with the double-stranded DnaA box, but also binds to the single-stranded DNA to promote unwinding. For example, DnaA protein can bind to single-stranded ATP-DnaA boxes mentioned above. The two-state and loop-back models can explain how DnaA protein melts DNA and stabilizes the unwound region by DnaA-ssDNA interaction [42]. In two-state model, DnaA protein guided from double-stranded DnaA boxes to the adjacent single-stranded DNA changes from a double-to a single-stranded binding mode. A new *oriC* element comprising repeated 3-mer motif (DnaA-trio), found in *Bacillus subtilis*, promotes DNA unwinding by stabilizing DnaA filaments on a single DNA strand [45]. Consequently, a basal unwinding system (BUS) comprised DnaA boxes and DnaA-trios in bacterial *oriC*s has been proposed [46]. Subsequent bioinformatic analyses of *oriC*s from over 2000 bacterial species, together with molecular biology studies of six representative species, found that the BUS is broadly conserved in bacteria [35]. Integration host factor (IHF) induces DNA to bend backwards in the loop-back model, bringing the DUE close to the DnaA protein bound to the DnaA box and thus simultaneously facilitating protein binding to double- and single-stranded DNA sequences. This mechanism has been identified in *E. coli* [47], and a similar mechanism might be also found in *Helicobacter pylori* [48] with a bipartite *oriC* and *V. cholerae* chromosome 2 whose replication initiator requires RctB protein other than DnaA protein [49].

In addition to binding sites for the DnaA protein, *oriC* has other binding sites for proteins that regulate replication initiation. Factor for inversion stimulation (Fis) and IHF bind to specific sites and bend *oriC* DNA to inhibit or facilitate DnaA binding in *E. coli* [47]. SeqA blocks *oriC* recognition of DnaA by binding to the transiently hemimethylated GATC sequence cluster [50]. The regulatory mechanisms might differ because of the diversity of regulatory proteins and their binding motifs among species. For example, CtrA in *Caulobacter crescentus* plays a similar role to SeqA and inhibit replication initiation by binding motifs (TTAA-N7-TTAA) [51, 52]. Wolanski *et al*. comprehensively summarized the detailed information about the proteins that regulate DNA replication initiation and their binding sites [53].

To facilitate a comprehensive understanding of the replication mechanism and sequence characteristics related to *oriC*s, Ori-Finder 2022 annotates various regulatory proteins and functional elements within *oriC*s. Updated information about the user interface, prediction framework, visualization, and analysis modules are described in detail below.

## Method

### Software implementation

Ori-Finder 2022 was deployed using a Linux-Apache-MySQL-PHP structure and mainly developed using Python and C++ languages. We packaged the pipeline into a container using Docker to ensure reproducible and reliable execution. We also integrated the third-party tools BLAST+ 2.11.0 [54], Prodigal [55], Stress-Induced Structural Transitions (SIST) [56], and MEME 5.4.1 [57], into Ori-Finder 2022 and tested the updated server on the web browsers, Firefox, Chrome, Safari, and Microsoft Edge.

### Input file

By December 28, 2021, 91.5% of 362,223 bacterial genomes in the NCBI Genome database were draft genomes with scaffold or contig assembly levels. We updated Ori-Finder to enable *oriC* prediction to meet the imperative need to annotate *oriC*s in these genomes (**Figure 1A**). The updated web server can consequently handle complete or draft bacterial genomes with or without annotations. Ori-Finder 2022 integrates the gene-finding algorithm Prodigal [55] to predict protein-coding genes in unannotated genomes in the FASTA format. If an annotated genome file is uploaded in the GBK format, the annotation information is automatically extracted by parsing text.

**Figure 1.**
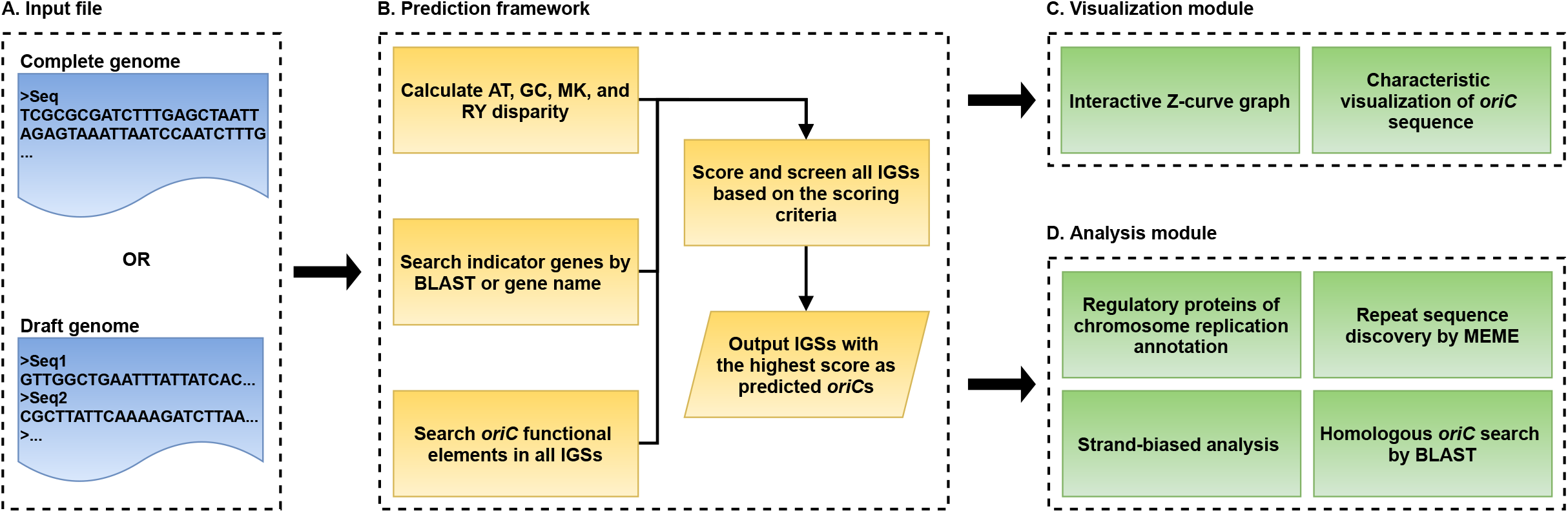
Workflow of Ori-Finder 2022. **A**. Input file of Ori-Finder 2022. Users can submit complete or draft genome in FASTA or GBK format. **B**. Prediction framework of Ori-Finder 2022. Ori-Finder 2022 predicts *oriC*s by comprehensively assessing DNA asymmetry, indicator genes, and *oriC* functional elements. **C**. Visualization module of Ori-Finder 2022. **D**. Analysis module of Ori-Finder 2022. *oriC*, replication origin; IGS, intergenic sequence.

### Updated prediction framework

Ori-Finder was originally developed with DNA asymmetry analysis using the Z-curve method, the distribution of DnaA boxes, and indicator genes close to *oriC*s [17]. Considering more *oriC* characteristics, the updated prediction framework of Ori-Finder 2022 adopts a new scoring criterion to quantitatively reflect these *oriC* characteristics of each intergenic sequence (IGS), and the IGSs with highest score are predicted as potential *oriC*s (**Figure 1B** and Table S1). As a characteristic of base composition, GC asymmetry is widely used for predicting *oriC*s. Ori-Finder 2022 scores the characteristics of base composition according to the distance to the minimum of the GC disparity (Table S1). Bacterial *oriC*s are usually adjacent to a *dnaA* gene, which can sever as an indicator for *oriC*s, but such genes are often different among bacterial species. Ori-Finder 2022 scores the characteristics of indicator genes, which can be adjusted based on the lineage and chromosome type entered by the user (Table S2). Ori-Finder 2022 scores DnaA boxes according to their numbers and mismatches. In addition, Ori-Finder 2022 identifies other functional elements of *oriC*, such as the Dam methylation site (GATC), and DnaA-trio, to screen prediction results if several IGSs with the same highest scores occur during the prediction process. For draft genomes, each sequence fragment will be predicted using Ori-Finder 2022, and all results will be considered together using the same prediction framework. Unlike the complete genome, the GC disparity minimum of each sequence fragment was used when scoring base composition.

### Updated user interface

According to the updated prediction framework, the user interface for data submission was redesigned to enhance user experience (**Figure 2A**). Ori-Finder 2022 only requires users to upload the genome file in FASTA or GBK format to deliver a default *oriC* prediction; moreover, it provides some customization parameters. In Ori-Finder 2022, the principal indicator gene is *dnaA* by default and will be adjusted according to the lineage and chromosome type entered by users (Table S2). The default DnaA box is the standard motif (TTATCCACA) of *E. coli*, while the built-in DnaA box motif can be selected according to the organism or lineage of the input genome. The drop-down checkboxes of the DnaA box motif and *dif* motif can achieve certain linkages for user convenience. Because of the diversity of DnaA boxes, Ori-Finder 2022 allows users to define their own DnaA box motifs. Users can select or define the *dif* motif in a similar way. Users can also choose strand-biased analysis for complete genomes.

**Figure 2.**
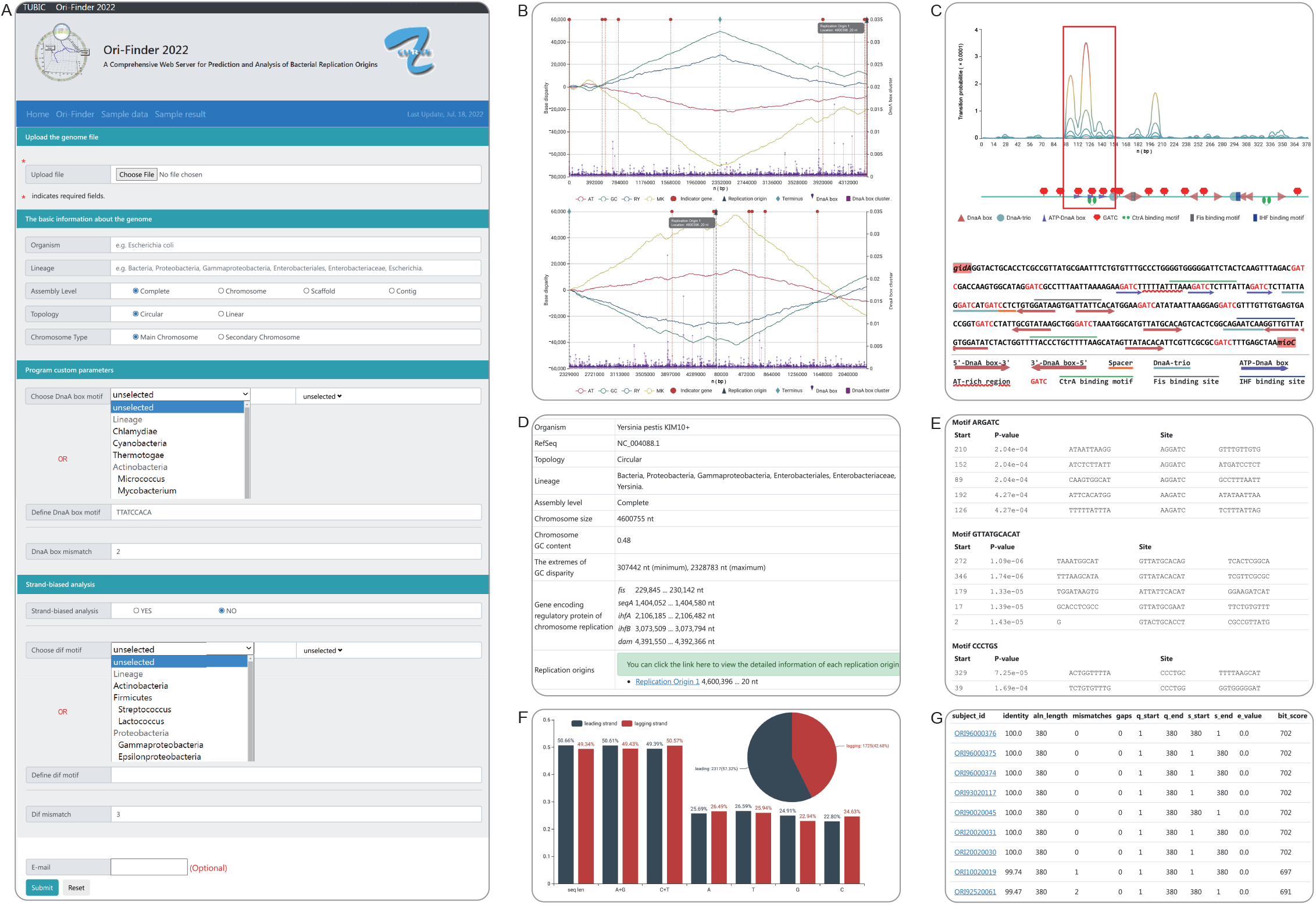
User interface and predicted result of Ori-Finder 2022 for *Yersinia pestis* KIM+. **A**. User interface of Ori-Finder 2022. **B**. Interactive Z-curve graph (original Z-curve graph and the one rotated at maximum of GC disparity) with *oriC* related information. **C**. Sequence characteristic visualization of the predicted *oriC*. **D**. Table in HTML providing basic information of *Y. pestis* KIM+ genome and its potential *oriC*. Genes encoding regulatory proteins of chromosome replication in the genome is provided here. **E**. Repeat sequence discovery by MEME. **F**. Strand-biased analysis. Pie and bar charts respectively show distribution of genes and bases in leading and lagging strands. **G**. Homologous *oriC* search by BLAST. *oriC*, replication origin.

### Updated visualization module

The updated visualization module in Ori-Finder 2022 contains interactive Z-curve graph and characteristic visualization of *oriC* sequences. Global or local information of the genome can be grasped at a glance from the interactive Z-curve graph that displays the four disparity curves and the distribution of DnaA boxes, indicator genes, potential *oriC*s, and replication terminus (**Figure 2B**). The red, green, blue, and yellow line graphs indicate the AT, GC, RY, and MK disparity curves, respectively, calculated according to the Z-curve method. The purple vertical line displays the density of DnaA boxes, which is used to indicate the existence of DnaA box cluster. Red, dark blue, and light blue dotted lines indicate locations of the indicator genes, *oriC*s, and replication terminus, respectively. The indicator genes were identified by parsing the annotation information of the genome or BLAST with known protein sequences encoding indicator genes. Ori-Finder 2022 can also predict the replication terminus of a complete genome according to the *dif* motif or the maximum of GC disparity. Users can select all the information or only several datasets to analyze according to their requirements. The graph also supports the zoom function for analyzing the details. Moreover, when users hover the cursor over the dotted lines marking predicted *oriC*s, indicator genes or replication terminus, the exact location and other information are automatically displayed.

The other visualization result provided by Ori-Finder 2022 is the characteristic visualization of *oriC* sequence, which displays the distribution of its functional elements. The first part is the line graph (**Figure 2C**, top), which shows the transition probability of each base pair in the sequences calculated using Stress-Induced Duplex Destabilization method [56] that analyzes stress-driven DNA strand separation. Five lines with gradient colors were calculated using different negative superhelicity values, and the peaks corresponded to the AT-rich sequence that might serve as a DUE. The second part is an *oriC* sequence schematic diagram showing the distribution of functional elements, such as DnaA box, DnaA-trio, ATP-DnaA box, GATC motif, and binding sites of CtrA, Fis, and IHF found in the predicted *oriC*s (Figure 2C, middle). The third part is the sequence of the predicted *oriC* in which the different elements are labeled with colors or symbols (Figure 2C, bottom). Indicator genes upstream and downstream of the predicted *oriC* are also labeled. In order to display the possible functional elements as comprehensively as possible, all possible DnaA trios are labeled, and a less conserved DnaA box with ≤ 4 mismatches from the standard DnaA box motif adjacent to a potential DnaA-trio will also be labeled, although its mismatch might be more than that entered by users.

### Updated analysis module

Ori-Finder 2022 was expanded to include the new analysis modules. Combined with the different elements labeled in *oriC* sequence (Figure 2C), the annotation of corresponding regulatory proteins, such as Fis, SeqA, and CtrA (**Figure 2D**), by Ori-Finder 2022 might provide new insights into the related regulatory mechanisms. In addition, the repeat sequences in predicted *oriC*s were discovered by MEME are listed in a HTML table to reveal possible new motifs (**Figure 2E**). A strand-biased analysis can reveal the distribution of genes and bases on the leading and lagging strands of a complete genome (**Figure 2F**). Sequences homologous to predicted *oriC*s were searched using BLAST against the DoriC database [31], and the BLAST results linked to the corresponding entry in the DoriC database are also provided (**Figure 2G**).

## Result and discussion

Here, *Yersinia pestis* KIM+ is presented to illustrate details of the predicted results of Ori-Finder 2022. The structure of *oriC* in *Y. pestis* KIM+ is similar to that in *E. coli* [58]. Figure 2 shows the main visualization and analytical results of the *oriC* predicted by Ori-Finder 2022 and the complete predicted results are available as a Sample result at our website (http://tubic.tju.edu.cn/Ori-Finder2022/public/index.php/retrieve/sample_result). Due to possible rearrangement, the four disparity curves of this genome fluctuate at their extrema [58], which does not seem to provide sufficient evidence to identify an *oriC* (Figure 2B). Ori-Finder 2022 identified an IGS of 380 base pairs as the potential *oriC* by taking more characteristics into consideration, such as indicator genes, DnaA box cluster, and other functional elements. Like that in *E. coli*, the predicted *oriC* in *Y. pestis* KIM+ was located between *gidA* and *mioC*. The sequence corresponding to the peak of the lines calculated by SIST also contained DnaA-trios and three ATP-DnaA boxes (AGATCT), which was likely to contain a site of DNA duplex unwinding (Figure 2C). The genome of *Y. pestis* KIM+ encodes regulatory proteins such as Fis, SeqA, IHF, and Dam (Figure 2D), and the possible binding sites for corresponding proteins are also found in the predicted *oriC*. Although the genome of *Y. pestis* KIM+ does not appear to encode CtrA proteins, two possible CtrA binding sites were identified within the predicted *oriC*. The repeat sequences in the predicted *oriC*s were discovered using MEME, which might reveal new *oriC* motifs. For example, two of the five motifs in the first set (ARGATC) overlapped with predicted ATP-DnaA boxes. In the second set (GTTATGCACAT), three of the five motifs overlapped with the predicted DnaA boxes, and the other two contained DnaA box-like motifs with three and four mismatches, respectively, from the perfect DnaA box (TTATCCACA) in *E*.*coli*. A *dif* site was located near the top of the GC disparity curve. Strand-biased analysis reveals the differences in some features between the leading and lagging strands. The lengths of the leading (50.66%) and lagging (49.34%) strands were almost identical. The leading strand included 2296 (56.76%) genes, probably a result of rearrangement during which the strand-biased phenomenon of genes is not obvious. The adenine (A), thymine (T), guanine (G), cytosine (C), purine (A+G), and pyrimidine (C+T) contents of the leading and lagging strands are also calculated (Figure 2F). The predicted result was considered reliable because homologous sequences were found in the DoriC database (Figure 2G).

## Conclusion

Ori-Finder has been widely applied by biologists over the past decade to predict bacterial *oriC*s, and some predictions have been experimentally confirmed [21-24] or supported by various studies [45, 59]. For example, the *oriC*s of 132 gut microbes in metagenomic samples predicted by metagenomic analyses and Ori-Finder were consistent (R^2^ = 0.98, P < 10^−30^) [59]. The bacterial *oriC* element, DnaA-trio, was found in 85% of *oriC*s predicted or confirmed from > 2000 species. Numerous bacterial *oriC*s generated by Ori-Finder have been used for large-scale data mining and analysis. Ori-Finder 2022 can now predict *oriC*s in complete or draft genomes based on an updated prediction framework and provides an interactive visualization module as well as a new analysis module. Ori-Finder will be continuously improved by incorporating state-of-the-art research results and integrating additional analysis modules. We plan to provide users with an integrated platform for comprehensive prediction, analysis, and knowledge mining to determine microbial replication origins. This will be achieved by integrating Ori-Finder 2 [60] that predicts archaeal replication origins, and Ori-Finder 3 [61], an online service for predicting replication origins in *Saccharomyces cerevisiae* in the future.

## Availability

Ori-Finder 2022 is freely available at:

http://tubic.tju.edu.cn/Ori-Finder2022 and https://tubic.org/Ori-Finder2022.

## CRediT author statement

**Meijing Dong:** Software, Writing-original draft, Visualization. **Hao Luo:** Software, Writing-original draft, Visualization. **Feng Gao:** Conceptualization, Writing-review & editing, Supervision, Funding acquisition. All authors read and approved the final manuscript.

## Acknowledgments

This work was supported by the National Key R&D Program of China (Grant No. 2018YFA0903700); and the National Natural Science Foundation of China [Grant Nos 21621004 and 31571358]. We thank Professor Chun-Ting Zhang for the invaluable assistance and inspiring discussions.

## Supplementary material

**Table S1 Scoring criterion of Ori-Finder 2022**

**Table S2 Indicator genes list for different chromosome type and lineage**

