## Supplemental Table 1 for "Ori-Finder 2022: A Comprehensive Web Server for Prediction and Analysis of Bacterial Replication Origins"

**Table S1** **Scoring criterion of Ori-Finder 2022**

| **Process** | **Characteristics** | **Score** | | |
| --- | --- | --- | --- | --- |
|  |  | **+1** | **+2** | **+3** |
| Prediction | Indicator gene | There is one other gene between *oriC* and the indicator gene. | The adjacent gene to *oriC* is a secondary indicator gene. | The adjacent gene to *oriC* is a principal indicator gene. |
|  | DnaA box cluster | DnaA box number ≥ 3  DnaA box mismatch ≤$M_{max}$^1^ | DnaA box number ≥ 3 DnaA box mismatch ≤ 1 | DnaA box number ≥ 3 DnaA box mismatch = 0 |
|  | Base composition | GC distance ≤ 5% | GC distance^2^ ≤ 1% |  |
| Screening^3^ | GATC motif^4^ | GATC motif number ≥ 8 |  |  |
|  | BUS^5^ | DnaA-trio score^5^ ≥ 15 |  |  |

*Note:*

1. If the maximum DnaA box mismatch ($M_{max}$) set by the user is less than 2, the score will be +2 ($M_{max}=1$) or +3 ($M_{max}=0$) if the identified DnaA box number is no less than 3.

2. GC distance represents the percentage of the distance between predicted *oriC*s and GC disparity minimum to genome size.

3. If several intergenic sequences with the same highest scores occur during the prediction process, they will be continue to be scored and screened.

4. GATC motif represents the Dam methylation site. The IGSs are scored based on the number of GATC motifs only when the genome has the gene encoding SeqA protein. Previous studies revealed that there are eight conserved GATCs in the *oriC* consensus sequence of the Enterobacteriacae [(Judith *et al.*, PNAS, 1983)](https://doi.org/10.1073/pnas.80.5.1164), so at least 8 GATCs are required here. Furthermore, the statistical results based on DoriC 6.5 showed that 97% of the genomes with SeqA had at least 8 GATCs in their *oriC*s.
5. DnaA-trio score is calculated based on the scoring matrix of DnaA-trio in our recent work [(Pelliciari *et al.*, Nucleic Acids Research, 2021)](https://academic.oup.com/nar/article/49/13/7525/6312737). The score of the DnaA-trios identified by experiments in this work are no less than 15, so the DnaA-trio score is required to be at least 15 here.

*oriC*, replication origin; BUS, basal unwinding system.
