## Supplemental Table 2 for "Ori-Finder 2022: A Comprehensive Web Server for Prediction and Analysis of Bacterial Replication Origins"

**Table S2 Indicator genes list for different chromosome type and lineage**

| **Chromosome type** | **Lineage** | **Indicator genes^1^** |
| --- | --- | --- |
| Main chromosome | Default | ***dnaA*** *dnaN rpmH gidA hemE mioC hemB* |
|  | Gammaproteobacteria | *dnaA dnaN rpmH* ***gidA*** *hemE mioC hemB* |
|  | Caulobacterales | *dnaA dnaN rpmH gidA* ***hemE*** *mioC hemB* |
|  | Rickettsiales | *dnaA dnaN rpmH gidA* ***hemE*** *mioC hemB* |
|  | Rhizobiales | *dnaA dnaN rpmH gidA* ***hemE*** *mioC hemB* |
|  | Cyannobacteria | *dnaA* ***dnaN*** *rpmH gidA hemE mioC hemB* |
|  | Chlamydia | *dnaA dnaN rpmH gidA hemE mioC* ***hemB*** |
| Secondary chromosome | Default | *repA repC parA parB dnaA* |

*Note:* 1. Genes in bold indicate principal indicator genes and others indicate secondary indicator genes.
